# Girdin is a component of the lateral polarity protein network restricting cell dissemination

**DOI:** 10.1101/733329

**Authors:** Cornélia Biehler, Li-Ting Wang, Myriam Sévigny, Alexandra Jetté, Clémence Gamblin, Rachel Catterall, Élise Houssin, Luke McCaffrey, Patrick Laprise

## Abstract

Epithelial cell polarity defects support cancer progression. It is thus crucial to decipher the functional interactions within the polarity protein network. Here we show that *Drosophila* Girdin and its human ortholog (GIRDIN) sustain the function of crucial lateral polarity proteins by inhibiting the apical kinase aPKC. Loss of GIRDIN expression is also associated with overgrowth of disorganized cell cysts. Moreover, we observed cell dissemination from *GIRDIN* knockdown cysts and tumorspheres, thereby showing that GIRDIN supports the cohesion of multicellular epithelial structures. Consistent with these observations, alteration of *GIRDIN* expression is associated with a poor overall survival in subtypes of breast and lung cancers. Overall, we discovered a core mechanism contributing to epithelial cell polarization from flies to humans. Our data also indicate that GIRDIN has the potential to impair the progression of epithelial cancers by preserving cell polarity and restricting cell dissemination.

## Introduction

The ability of epithelia to form physical barriers is provided by specialized cell-cell junctions, including the *zonula adherens* (ZA). The latter is a belt-like adherens junction composed primarily of the transmembrane homotypic receptor E-cadherin, which is linked indirectly to circumferential F-actin bundles through adaptor proteins such as β-catenin and α-catenin (Desai et al., 2013; Harris and Tepass, 2010). In *Drosophila* embryonic epithelia, the protein Girdin stabilizes the ZA by reinforcing the association of the cadherin–catenin complex with the actin cytoskeleton (Houssin et al., 2015). This function in cell–cell adhesion is preserved in mammals, and supports collective cell migration (Wang et al., 2018; Wang et al., 2011). Fly and human Girdin also contribute to the coordinated movement of epithelial cells through the organization of supracellular actin cables (Houssin et al., 2015; Wang et al., 2018).

In addition to creating barriers, epithelial tissues generate vectorial transport and spatially oriented secretion. The unidirectional nature of these functions requires the polarization of epithelial cells along the apical-basal axis. In *Drosophila*, the scaffold protein Bazooka (Baz) is crucial to the early steps of epithelial cell polarization, and for proper assembly of the ZA (Bilder et al., 2003; Harris and Peifer, 2004; McGill et al., 2009; Muller and Wieschaus, 1996). Baz recruits atypical Protein Kinase C (aPKC) together with its regulator Partitioning defective protein 6 (Par-6) to the apical membrane (Harris and Peifer, 2005; Tepass, 2012; Wodarz et al., 2000). Baz also contributes to apical positioning of the Crumbs (Crb) complex, which is composed mainly of Crb, Stardust (Sdt), and PALS1-associated Tight Junction protein (Patj) (Krahn et al., 2010; Tepass, 2012). Once properly localized, the aPKC–Par-6 and Crb complexes promote the apical exclusion of Baz, which is then restricted to the ZA (Harris and Peifer, 2005; Morais-de-Sa et al., 2010; Walther and Pichaud, 2010). The apical exclusion of Baz is essential to the positioning of the ZA along the apical-basal axis (Morais-de-Sa et al., 2010), and for full aPKC activation (David et al., 2013; McCaffrey et al., 2012; Soriano et al., 2016).

The function of aPKC is evolutionarily preserved, and human atypical PKCι and PKCζ (PKCλ in mammals) contribute to epithelial cell polarization (Hong, 2018). aPKC maintains the identity of the apical domain through phospho-dependent exclusion of lateral polarity proteins such as Yurt (Yrt) and Lethal (2) giant larvae (Lgl) (Gamblin et al., 2014; Gamblin et al., 2018; Hutterer et al., 2004; Laprise and Tepass, 2011). In return, these proteins antagonize the Crb- and aPKC-containing apical machinery to prevent the spread of apical characteristics to the lateral domain (Bilder et al., 2003; Drummond and Prehoda, 2016; Fletcher et al., 2012; Gamblin et al., 2014; Hutterer et al., 2004; Laprise et al., 2006; Laprise et al., 2009; Tanentzapf and Tepass, 2003; Yamanaka et al., 2006). In combination with the function of Baz, these feedback mechanisms provide a fine-tuning of aPKC activity in addition to specifying its subcellular localization. This is crucial, as both over- and under-activation of aPKC is deleterious to epithelial polarity in fly and mammalian cells, and ectopic activation of aPKC can lead to cell transformation (Archibald et al., 2015; David et al., 2010; Lin et al., 2000; Sotillos et al., 2004; Wodarz et al., 2000).

Cell culture work has established that mammalian GIRDIN interacts physically with PAR3 – the ortholog of Baz – and PKCλ (Goldstein and Macara, 2007; Ohara et al., 2012; Sasaki et al., 2015). Depletion of *GIRDIN* in Madin-Darby Canine Kidney (MDCK) epithelial cells delays the formation of tight junctions in Ca^2+^ switch experiments (Sasaki et al., 2015). GIRDIN is also an effector of AMP-activated protein kinase (AMPK) in the maintenance of tight junction integrity under energetic stress (Aznar et al., 2016b). Moreover, mammalian GIRDIN is required for the formation of epithelial cell cysts with a single lumen, supporting a role for this protein in epithelial morphogenesis as reported in flies (Aznar et al., 2016b; Houssin et al., 2015; Ohara et al., 2012; Sasaki et al., 2015). As cyst morphogenesis is linked to epithelial cell polarity (Sigurbjornsdottir et al., 2014), these studies suggest that GIRDIN is involved in establishing the apical-basal axis. However, further studies are required to clarify the role of GIRDIN in apical-basal polarity *per se*, as other cellular processes could explain the phenotype associated with altered GIRDIN expression. For instance, spindle orientation defects impair the formation of epithelial cysts (Jaffe et al., 2008). Of note, PAR3, aPKC, and AMPK are all required for proper spindle positioning in dividing epithelial cells (Durgan et al., 2011; Hao et al., 2010; Lu and Johnston, 2013; Thaiparambil et al., 2012). The molecular mechanisms sustaining the putative role of GIRDIN in epithelial cell polarity also need to be better deciphered. Here, we further investigated the role of fly and human Girdin proteins in the regulation of epithelial cell polarity, and showed that these proteins are part of the lateral polarity protein network. One crucial function of Girdin proteins is to repress aPKC function. We also discovered that loss of Girdin proteins promotes overgrowth of cell cysts, and cell dissemination from these multicellular structures. Consistent with these data, we found that low *GIRDIN* expression correlates with poor overall survival in subtypes of breast and lung cancers.

## Results

### *Girdin* is a crucial component of the genetic network supporting lateral membrane stability in polarized epithelial cells

To explore the role of Girdin in epithelial cell polarity regulation, we investigated its functional relationship with Yrt and Lgl, which are known polarity regulators in *Drosophila* embryos. These lateral proteins prevent overactivation of the Crb- and aPKC-containing apical machinery, thereby precluding apicalization of the lateral membrane (Bilder et al., 2003; Gamblin et al., 2014; Laprise et al., 2006; Tanentzapf and Tepass, 2003; Yamanaka et al., 2006). Similar to Girdin, Yrt and Lgl are provided maternally (Bilder, 2004; Houssin et al., 2015; Laprise et al., 2006). Although suboptimal, the maternal contribution is sufficient to maintain apical-basal polarity in zygotic mutant embryos carrying null alleles for *Girdin*, *yrt* or *lgl* (Bilder, 2004; Houssin et al., 2015; Laprise et al., 2006) (Figure 1A-I, and 2A-C). Zygotic loss of these genes thus represents a sensitized background. In contrast, a complete loss of Lgl expression in *lgl* maternal and zygotic (M/Z) mutants is associated with strong polarity defects characterized by ectopic localization of the apical protein Crb to the lateral membrane in post-gastrulating embryos (Tanentzapf and Tepass, 2003). This phenotype is clearly visible at stage 11 of embryogenesis as shown by the partial co-localization of Crb with the lateral protein Discs large 1 (Dlg1; Figure 1J). At stage 13, the monolayered architecture of the differentiating epidermis is compromised, and polarity defects were attenuated but still apparent (Figure 1K). Toward the end of embryogenesis (stage 16), Lgl-deficient epidermal cells formed polarized cysts of cells with their extended – cuticle secreting – apical membrane facing out (Figure 1L). As a consequence, *lgl* M/Z mutant embryos assembled a highly convoluted cuticle (Figure 1R), whereas zygotic *lgl* and *Girdin* mutant embryos displayed a normal cuticle (Figure 1P, Q). Strikingly, a combination of zygotic *lgl* and *Girdin* mutations phenocopied a total loss of Lgl (Figure 1M-O, S). Similarly, depletion of Girdin, which is not sufficient to cause obvious polarity defects (Houssin et al., 2015) (Figure 2D-F), strongly enhanced the zygotic *yrt* mutant phenotype. Indeed, *Girdin yrt* double mutant specimens were indistinguishable from *yrt* M/Z embryos (Laprise et al., 2006) (Figure 2G-L, and O, P), and presented an expansion of Crb expression territories at stages 13 of embryogenesis (Figure 2H, K). These strong genetic interactions suggest that Girdin, Yrt, and Lgl share a common function in maintaining lateral characteristics in polarized epithelial cells.

**Figure 1.**
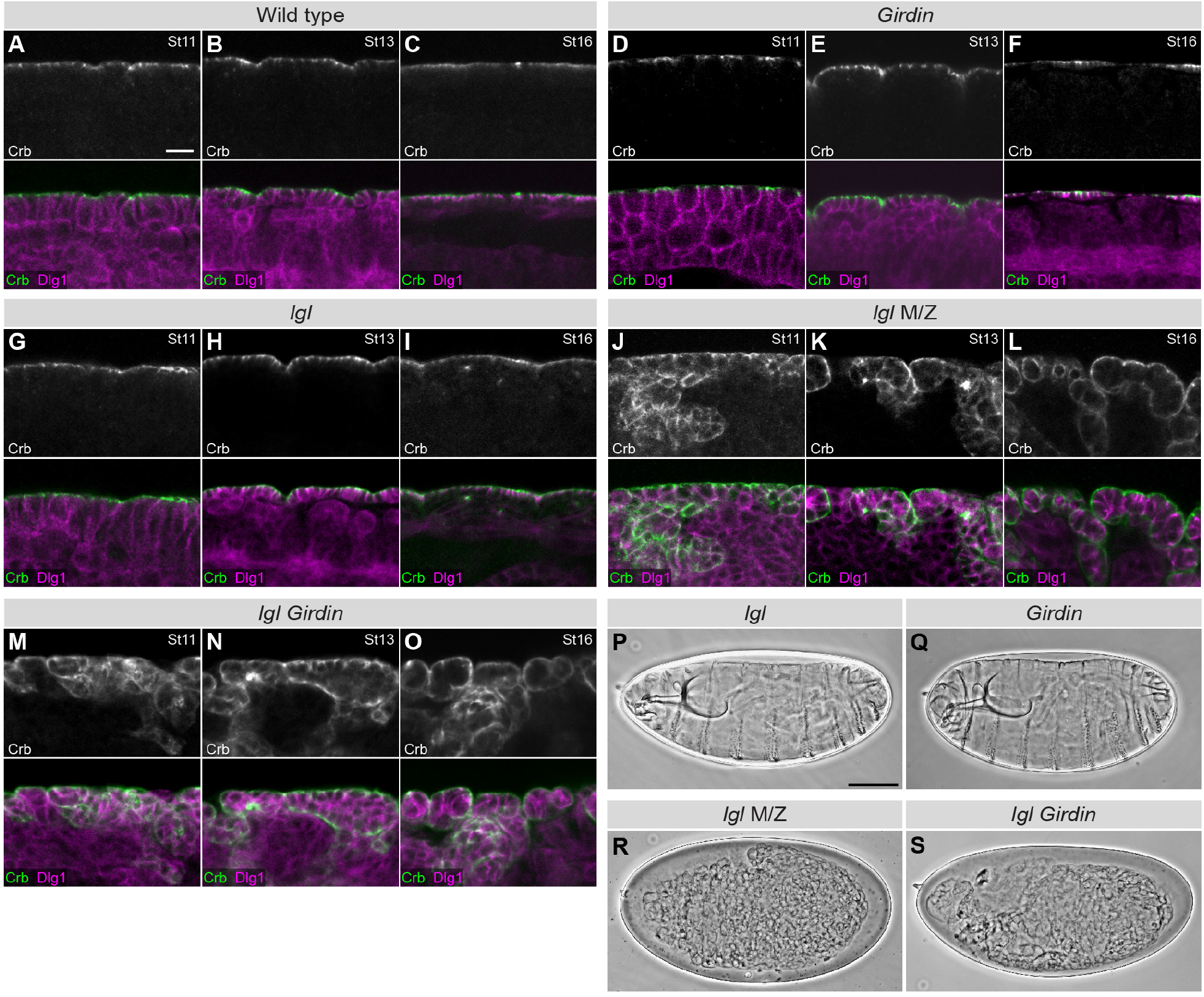
Girdin cooperates with Lgl to support epithelial cell polarity. **A-O**, Embryos of the indicated genotypes were fixed and co-stained for Crb and Dlg1. M/Z stands for maternal and zygotic mutant. Panels depict a portion of the ventral ectoderm or epidermis at stage (St) 11 (**A**, **D**, **G**, **J**, **M**), stage 13 (**B**, **E**, **H**, **K**, **N**), or stage 16 (**C**, **F**, **I**, **L**, **O**) of embryogenesis. In each case, the apical membrane is facing up. Scale bar in **A** represents 10 μm, and also applies to **B**-**O**. **P**-**S**, Panels depict cuticle preparations of whole mounted embryos of the indicated genotypes. The anterior part of the embryo is oriented to the left, and the dorsal side is facing up. Scale bar in **P** represents 100 μm, and also applies to **Q**-**S**.

**Figure 2.**
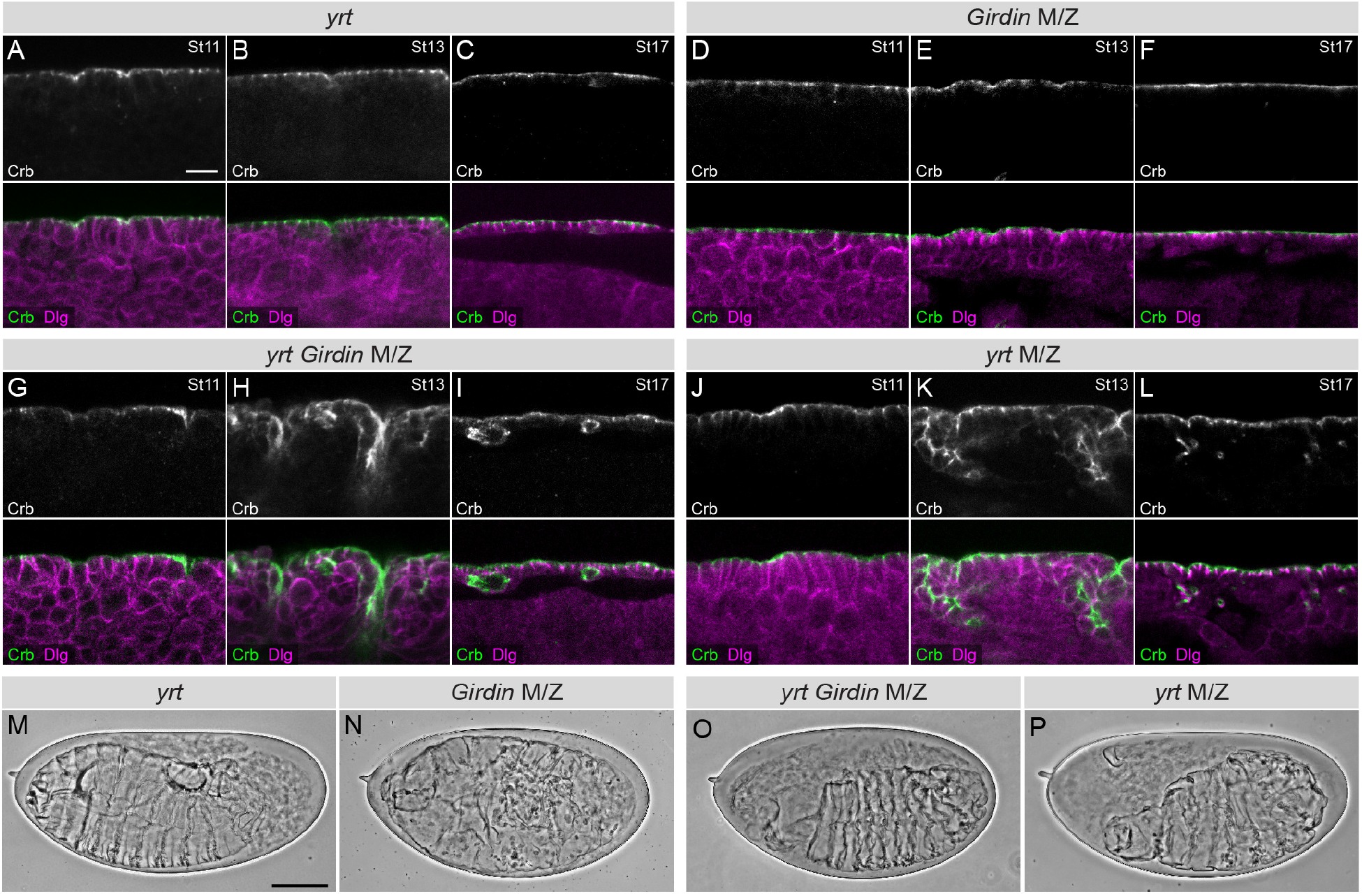
*Girdin* mutation enhances the delocalization of the apical protein Crb in *yrt* mutants. **A-L**, Embryos were immunostained for Crb and Dlg1, and a part of the ventral ectoderm or epidermis was imaged by confocal microscopy (the apical side of the epithelial tissue is facing up). The developmental stage of embryos (St) is indicated in the upper right corner of panels. Scale bar in **a** represents 10 μm, and also applies to **B**-**L**. **M**-**P**, Cuticle preparations of whole mounted embryos are shown. Scale bar in **M** represents 100 μm, and also applies to **N**-**P**.

To clarify the function of Girdin within the lateral polarity network, we performed epistasis analysis by combining the *Girdin* mutant allele with a null (M/Z) background for *yrt*. The severity of Crb delocalization to the lateral membrane was similar in *yrt* M/Z embryos and in *yrt* M/Z *Girdin* double mutant embryos, and no additional phenotypes were apparent at the cellular level or at the level of tissue organization (Figure 2J-L, P, and 3D-F, N). This shows that *yrt* is epistatic to *Girdin*. Thus, these genes act in a common genetic pathway in which *yrt* acts downstream of *Girdin*. We tried to carry out similar experiments with *lgl*, but we faced technical challenges and were not able to obtain *lgl* M/Z *Girdin* specimens. As an alternative, we used a maternal knockdown (kd) approach, which closely mimicked a complete loss of Lgl (Figure 1J-L, R and 3G-I, O; Supplementary Figure 1A). *lgl* (kd) *Girdin* embryos were similar to their *lgl* (kd) counterparts (Figure 3G-L, O-P). Our results thus suggest that *lgl* is epistatic to *Girdin*, and that *lgl* is downstream of *Girdin* in this newly defined pathway. Taken together, our data argue that Girdin is an important upstream positive regulator of both Yrt and Lgl function.

**Figure 3.**
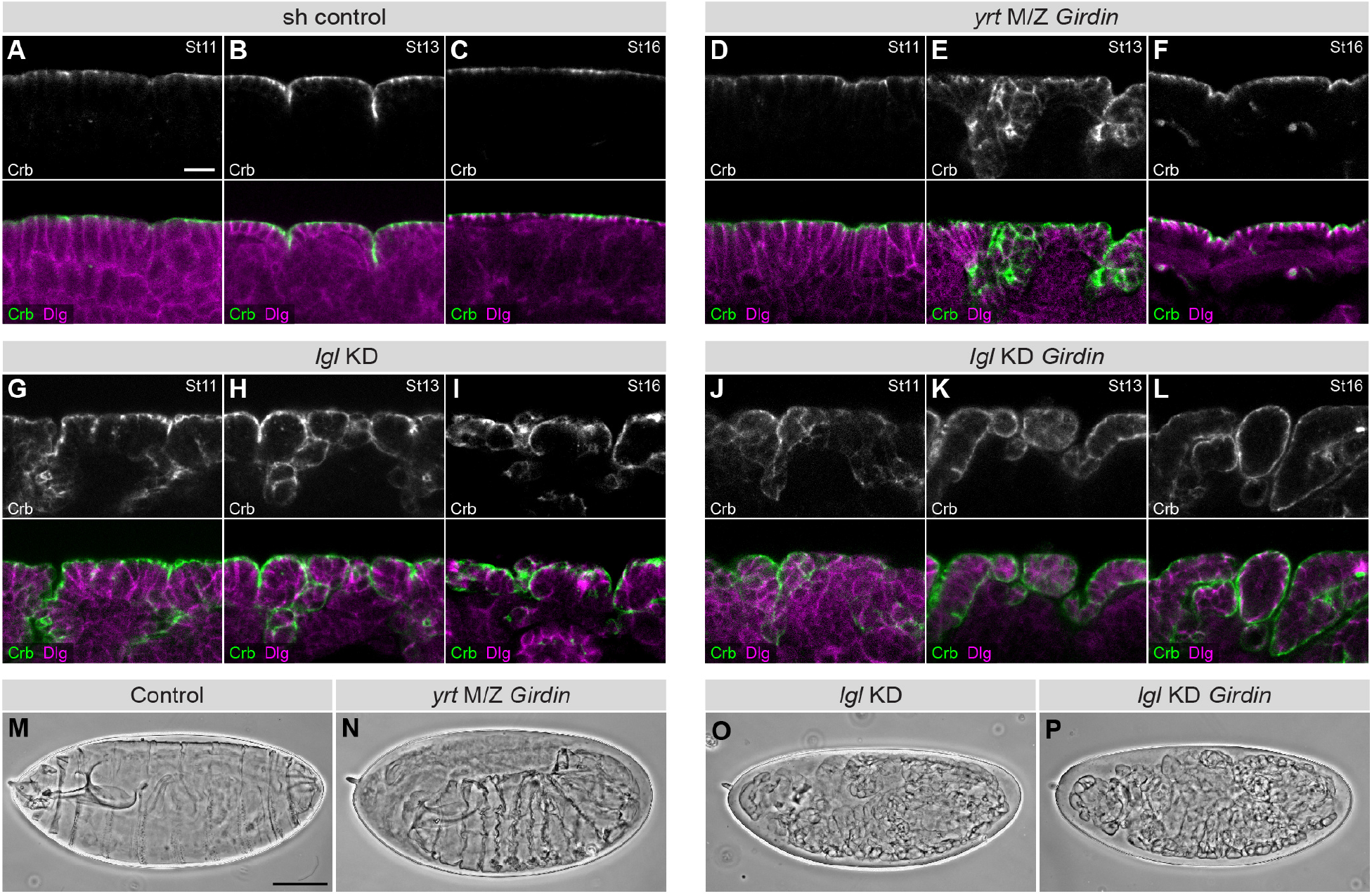
Girdin is part of the lateral polarity protein network. **A-L**, Immonostaining of Crb and Dlg1 in embryos of the indicated genotypes are shown. Panels show a portion of the ventral side of embryos at different stages (St) of development. Upper panels depict Crb staining, whereas lower panels display the merged images of Crb and Dlg1 staining. Scale bar in **A** represents 10 μm, and also applies to **B**-**L**. **M**-**P**, Panels show the cuticle secreted by embryos of the indicated genotypes. The anterior side of embryos is on the right, and dorsal side is up. Scale bar in **M** represents 100 μm, and also applies to **N**-**P**.

### Fly Girdin and human GIRDIN repress the function of aPKC to support epithelial cell polarity

Although Yrt and Lgl control Crb activity independently and in separate time windows during fly embryogenesis (Laprise et al., 2006; Laprise et al., 2009; Tanentzapf and Tepass, 2003), they are both inhibited by aPKC-dependent phosphorylation (Bailey and Prehoda, 2015; Betschinger et al., 2003; Gamblin et al., 2014; Gamblin et al., 2018). It is thus plausible that Girdin acts as a repressor of aPKC activity to support the function of lateral proteins. Accordingly, Yrt, which is a substrate of aPKC (Gamblin et al., 2014), showed reduced electrophoretic mobility in *Girdin* null embryos compared to control animals. This was due to enhanced phosphorylation of Yrt, as treatment of samples with the λ Protein Phosphatase abolished the delayed migration profile of Yrt in *Girdin* null embryos (Figure 4A). This result strongly argues that the activity of aPKC increases in the absence of Girdin. To obtain functional evidence that Girdin represses aPKC activity, we performed genetic interactions. Maternal knockdown of *aPKC* resulted in strong epithelial morphogenesis defects as revealed by analysis of the cuticle. Indeed, most knocked-down embryos displayed epithelial tissue collapse (referred to as class I embryos, Figure 4B, E). Remaining embryos showed a weaker phenotype, and secreted either small patches of cuticle (class II; Figure 4C, E), or large continuous sheets of cuticle with differentiated structure such as denticle belts (class III; Figure 4D, E). Knockdown of *aPKC* in zygotic *Girdin* mutants, which show a normal cuticle phenotype (Houssin et al., 2015) (Figure 1Q), significantly changed the distribution of embryos in each category. Specifically, a decrease in class I embryos concomitant with an increase in class II and III specimens was observed (Figure 4E). This shows that reduction of Girdin levels alleviates the impact of *aPKC* knockdown, and suggests that Girdin normally limits the function of residual aPKC in knocked-down embryos (Supplementary Figure 1B). In accordance with this conclusion, overexpression of FLAG-Girdin enhanced the severity of the phenotype associated with the knockdown of *aPKC* (Figure 4F). Thus, *Drosophila* Girdin opposes aPKC function, and supports apical-basal polarity in epithelial cells.

**Figure 4.**
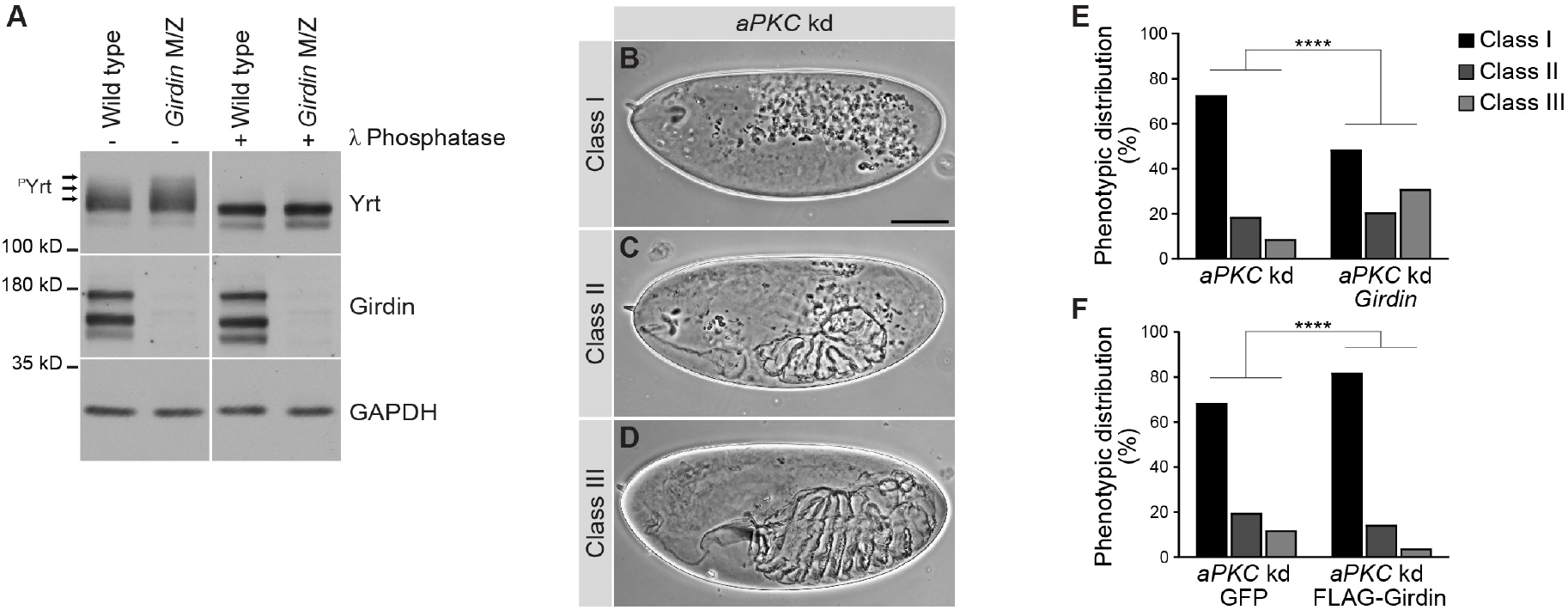
Girdin antagonizes aPKC functions. **A**, Stage 11–13 wild type embryos or maternal and zygotic (M/Z) *Girdin* mutant embryos were homogenized and processed for Western blotting. Where indicated, samples were treated with the λ Protein Phosphatase prior to electrophoresis. Arrows indicate the position of phosphorylated Yrt proteins (^P^Yrt). Immunoblotting of Girdin validates the genotype of embryos, whereas Gapdh1 was used as loading control. **B**-**D**, Cuticle preparations of *aPKC* knockdown (kd) embryos. Embryos were separated in three classes (I, II, or III) to account for phenotypic variability. Scale bar in **B** represents 100 μm, and also applies to **C**, **D**. **E**, Histogram showing the phenotypic distribution in percentage (%; according to the classes defined in **B**-**D**) in collections of embryos of the following genotypes: *aPKC* knockdown in a wild type background (*aPKC* kd; *n* = 1312), *aPKC* knockdown in a *Girdin* mutant background (*aPKC* kd *Girdin; n* = 1573). **F**, Phenotypic distribution of *aPKC* knockdown embryos expressing GFP (*aPKC* kd GFP; *n* = 603) or FLAG-Girdin (*aPKC* kd FLAG-Girdin; *n* = 697). For **E** and **F**, embryos were analyzed in three blinded experiments, and a Chi-squared test of independence was performed (**** : ρ < 0.0001).

To explore the intriguing possibility that human GIRDIN also restricts aPKC activity to control polarity, we used the human colon carcinoma Caco-2 cell line. These cells adopt a monolayered organization in 3D culture, and form hollow cysts with a single lumen. Cells forming the cysts are polarized along the apical-basal axis, with the PAR6- and F-ACTIN-enriched apical domain facing the central lumen [Figure 5A, H, L, N; (Jaffe et al., 2008)]. These proteins segregate from the lateral adhesion protein E-CADHERIN (E-CAD; Figure 5H-L and not shown). Consistent with our hypothesis, knockdown of *GIRDIN* in Caco-2 cells mimicked the phenotype associated with increased PKCζ activity (expression of the constitutively active PKCζ^T410E^), and resulted in the formation of solid cysts, or cysts with multiple atrophic lumens (Figure 5A,B, E, F and Supplementary Figure 1C). *GIRDIN* knockdown cells forming the disorganized 3D structures were mispolarized, as shown by the peripheral accumulation of PAR6 (Figure 5I, M, arrow) and F-ACTIN (Figure 5O). Despite that *GIRDIN* knockdown cysts displayed a similar size to controls, they contained more cells resulting from lumen filling (Figure 5N-P). The overgrowth phenotype was exacerbated and resulted in larger cysts when *GIRDIN* knockdown was combined with overexpression of wild type PKCζ, whereas overexpression of the latter alone had no impact on cyst size (Figure 5C-D, G). Again, this is consistent with a stronger PKCζ activation in *GIRDIN* knocked-down cells compared to cells expressing a scrambled shRNA, as expression of PKCζ^T410E^ also caused enlargement of cysts (Figure 5E, G). To confirm that aPKC activation contributes to the phenotypes associated with depletion of GIRDIN, we used the aPKC inhibitor CRT-006-68-54. Strikingly, inhibition of aPKC activity suppressed the defects caused by knockdown of *GIRDIN* (Figure 5Q-U). Together, these results indicate that, similar to fly Girdin, GIRDIN restrains aPKC function to support epithelial morphogenesis and apical-basal polarity.

**Figure 5.**
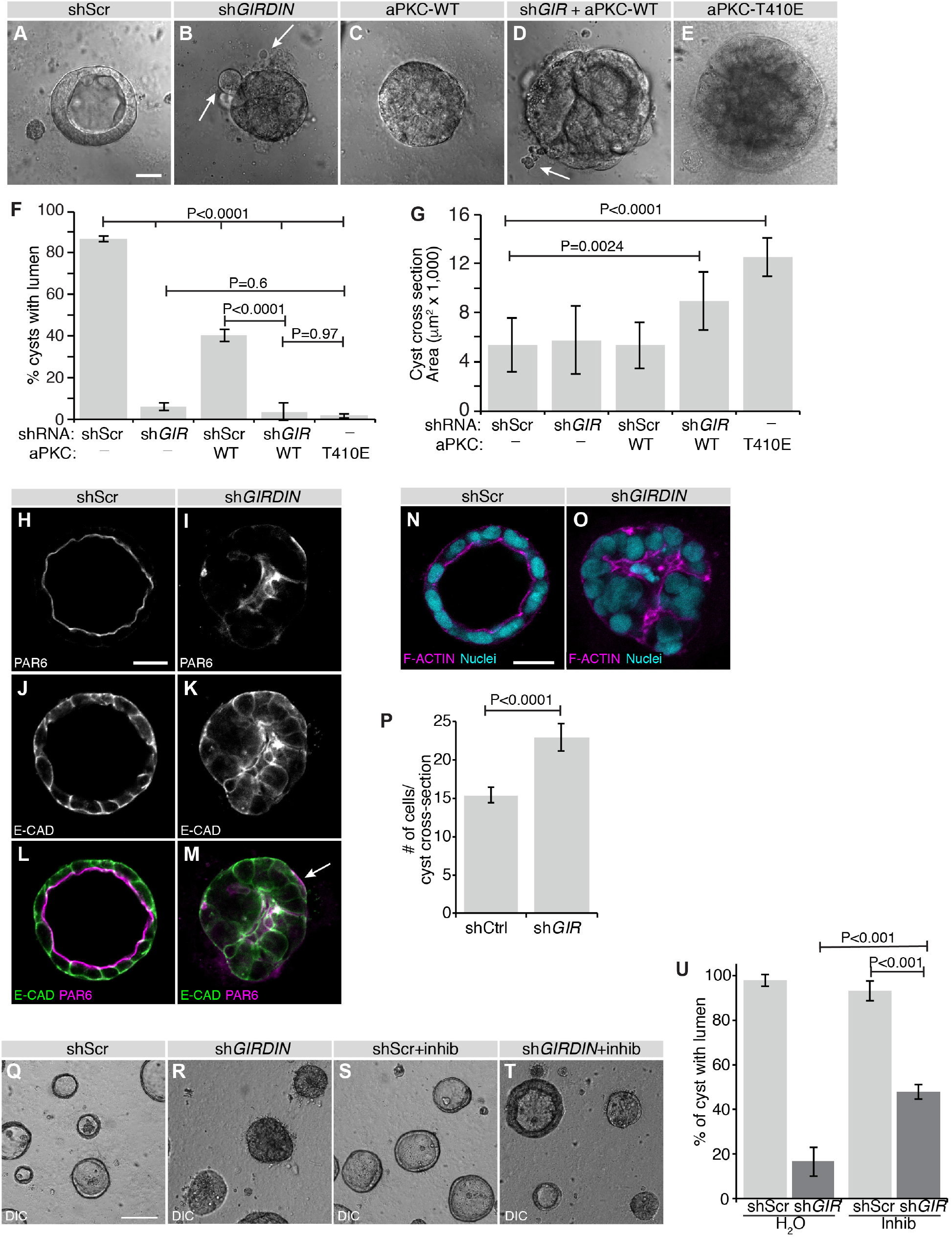
Human GIRDIN is essential for epithelial morphogenesis and polarity. **A-E**, Caco-2 cysts after 7 days in culture were observed by DIC microscopy, and the proportion of 3D cellular structures with a single prominent lumen was assessed (shScr, n=27; sh*GIR*, n=39 from 3 independent experiments). Arrows in **B** and **D** highlight dissociated cells. Scale bar represents 25 μm. **F**, Histogram displaying the luminal phenotypes observed in a-e. Error bars = sd. **G**, Histogram displaying mean sizes (cross-sectional area) of cysts. Error bars = sd. **H-M**, Caco-2 cysts after 7 days in culture were immunostained for PAR6 and E-CAD and visualized by confocal microscopy. Arrows show an example of a structure with Par6 localized basally. Scale bar represents 25 μm. **N-O**, Caco-2 cysts after 7 days in culture were stained with phalloidin (F-ACTIN) and Hoechst (nuclei) and visualized by confocal microscopy. Scale bar represents 25 μm. **P**, Histogram displaying the number of cells counted in cross-sectional images through the middle of 3D cellular aggregates. Error bars = sd. **Q-T**, Caco-2 cysts after 7 days in culture were visualized by DIC microscopy. Cells were treated with 1.5 μM of the aPKC inhibitor CRT-006-68-54 (inhib) or vehicle control (water) for the duration of the 3D culture period. Scale bar represents 100 μm. **U**, Histogram displaying the proportion of 3D cellular structures with a single prominent lumen (shScr, n=278; shGIR, n=181; shScr+Inhib, n=321; shGIR+Inhib, n=196 from 3 independent experiments). Error bars = sd. Differences were determined using ANOVA with Tukey HSD (**F**, **G**, **U**) or Student’s t-test (**P**).

### GIRDIN maintains the cohesion of epithelial cysts

In addition to the morphogenesis and polarity defects associated with decreased GIRDIN expression, we observed the presence of isolated cells or small cell aggregates in the proximity of most *GIRDIN* knocked-down Caco-2 cell cysts (Figure 5B and D, arrows). As we previously reported that cell cysts detach from the epidermis and survive outside of it in *Girdin* mutant *Drosophila* embryos (Houssin et al., 2015), we hypothesized that some cells separate from *GIRDIN* knockdown Caco-2 cysts. To test this hypothesis, we performed time-lapse microscopy of control (expressing a scrambled shRNA) and *GIRDIN* knocked-down cysts. Over a period of 26 hours, control cysts showed cell–cell rearrangements, which were resolved to maintain the monolayered organization (Figure 6A and video 1). In *GIRDIN* knockdown cysts, cells were frequently observed extending from the periphery of cysts and detaching (Figure 6B arrow, E-G, and video 2). In control cysts, the extensions were less frequent, and detachment was not observed (Figure 6A, E-G, and video 1). Further analysis also revealed that loss of GIRDIN expression is associated with budding from a large group of cells from the cyst (Figure 6C and video 3), or the fragmentation of cysts into multiple smaller cell aggregates (Figure 6D, video 4). Staining for viability revealed that all cells were alive prior to detachment from cysts, and some cell survived outside of their cyst of origin (Figure 6H-N). Expression of oncogenic KRAS (KRAS^G12V^), which is an important driver of colon cancer (Hardiman, 2018), improved the survival of detached cells (Figure 6O-S). Collectively, our results show that GIRDIN is required to maintain the cohesion of multicellular epithelial structures, and that its loss is associated with cell dispersion in control and cancer-mimetic contexts. This suggests that reduced GIRDIN may confer a more aggressive phenotype to cancer cells, and sustain tumor progression.

**Figure 6.**
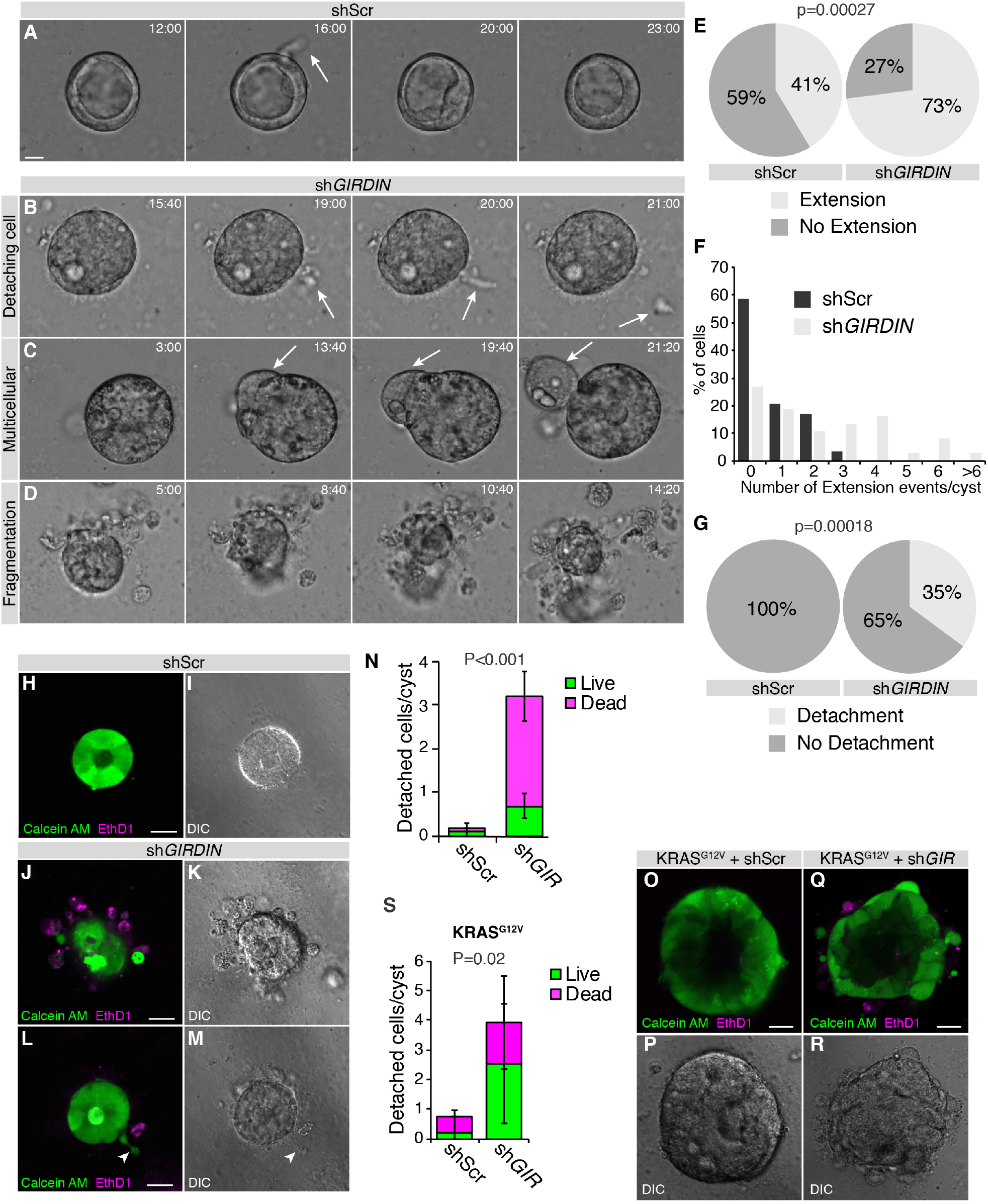
Human GIRDIN maintains cell polarity and prevents cell dissemination. **A**-**D**, Caco-2 cells were cultured for 7 days, then observed by time-lapse confocal microscopy every 20 min for 26 h. Still frames from videos display representative events observed (shScr, n=19; shGIR, n=28). Scale bar represents 25 μm. **E**, Proportion of control (shScr) and *GIRDIN*-knockdown cysts displaying transient cellular extensions. **F**, Distribution of the number of transient cellular extensions per cyst. **G**, Proportion of control (shScr) and *GIRDIN*-knockdown cysts from which cells disseminated. **H**-**M**, *GIRDIN* knockdown and control (shScr) Caco-2 cells were cultured for 7 days, then stained with calcein AM and ethidium homodimer to visualize live (green) and dead (magenta) cells by confocal microscopy. Scale bars represent 25 μm. **N**, Histogram displaying the mean number of cells detached from Caco-2 aggregates (shScr, n=27; shGIR, n=30) and the proportion of live and dead detached cells. **O**-**R**, *GIRDIN* knockdown (n=27) and control (shScr, n=27) KRAS^G12V^-expressing Caco-2 cells were stained with calcein AM and ethidium homodimer to visualize live (green) and dead (magenta) cells by confocal microscopy. Scale bars represent 25 μm. **S**, Histogram displaying the mean number of cells detached from cyst and the proportion of live and dead detached cells. Differences were determined using Chi-squared test (**E**, **G**) or ANOVA with Tukey HSD (**N**, **S**).

### Alteration of *GIRDIN* expression is associated with a poor prognosis in a subset of breast and lung cancers

To further investigate the relevance of our data to human cancer, we examined *GIRDIN* expression and patient survival outcomes. We observed that low *GIRDIN* expression was associated with reduced overall survival in more aggressive breast cancer types (Luminal B, HER2+, and basal Breast Cancer), compared to less aggressive Luminal A subtype (Figure 7A-D). Low *GIRDIN* was also associated with poorer survival in lung adenocarcinoma, but not lung squamous cell carcinoma (Figure 7E, F). Therefore, *GIRDIN* expression is linked to survival outcomes, and displays specificity for certain cancer types.

**Figure 7.**
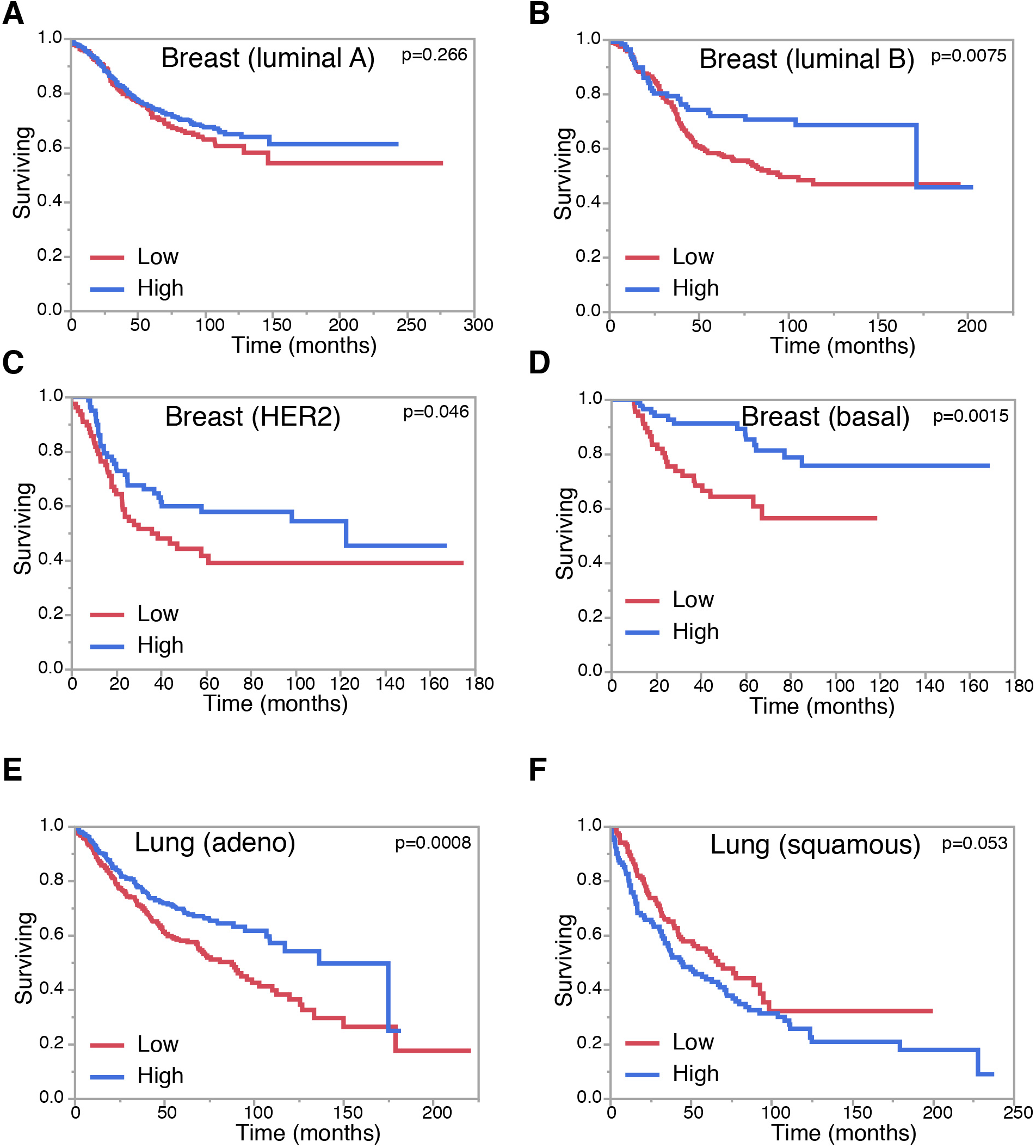
Altered *GIRDIN* expression correlates with survival in epithelial cancers. **A-G**, Kaplan-Meier survival plots for *GIRDIN* expression in breast cancer subtypes [luminal A (**A**), Luminal B (**B**), HER2-positive (**C**), and basal (**D**)], and lung cancer subtypes [adenocarcinoma (**E**), squamous (**F**)]. Differences between groups were determined using a Log-Rank test.

## Discussion

Using classical genetics in flies, we have shown that *Girdin* is part of the lateral polarity network. Mutation of *Girdin* exacerbates the polarity defects in zygotic *lgl* or *yrt* mutant embryos. It was previously proposed that maintenance of the ZA is essential to prevent the spread of apical characteristics to the lateral domain in *lgl* zygotic mutant embryos (Kaplan et al., 2009). Girdin is important in strengthening DE-cadherin (DE-cad)-dependent cell–cell adhesion in developing embryos (Houssin et al., 2015), suggesting that alteration of ZA properties could explain the genetic interaction between *Girdin* and *lgl*. However, we observed no genetic interaction between genes coding for core components of the ZA and *lgl* (not shown). We thus conclude that the weakening of cell-cell adhesion is not sufficient to enhance the *lgl* mutant phenotype, and that Girdin contributes directly to the protein network supporting the stability of the lateral domain. Specifically, we found that Girdin opposes the function of aPKC, which plays a crucial role in the establishment and maintenance of the apical domain (Hong, 2018; Tepass, 2012).

Our work demonstrates that the role of Girdin in restricting aPKC activity is evolutionarily conserved. This function confers on human GIRDIN the ability to maintain apical-basal polarity in Caco-2 cells, and to support epithelial cyst morphogenesis. These results are in line with previous studies suggesting a role for GIRDIN in polarity and cystogenesis in MDCK and MCF10A epithelial cells (Ohara et al., 2012; Sasaki et al., 2015). While aPKC has no impact on Girdin/GIRDIN expression in fly embryos and Caco-2 cells (not shown), PKCλ enhances GIRDIN expression in MDCK cells (Sasaki et al., 2015). Moreover, knockdown of *aPKC* or *GIRDIN* gives a similar phenotype characterized by defects in tight junction integrity and cyst formation (Nagai-Tamai et al., 2002; Sasaki et al., 2015; Suzuki and Ohno, 2006; Suzuki et al., 2001). It was thus proposed that GIRDIN is an effector of PKCλ (Sasaki et al., 2015). Although cell-type-specific mechanisms may exist, our data suggest that this hypothesis needs to be revisited in favor of a model in which the induction of GIRDIN expression by PKCλ in MDCK cells initiates a negative feedback loop instead of cooperation between these proteins. The fact that both overactivation of aPKC or inhibition of its activity is deleterious to epithelial cell polarity and cyst morphogenesis may underlie the conflicting interpretations of the data in the literature (Archibald et al., 2015; Suzuki and Ohno, 2006). GIRDIN is also known to modulate heterotrimeric G protein signaling (Aznar et al., 2016a; Ghosh et al., 2017) – a role that seems to contribute to the formation of normal cysts by MDCK cells (Sasaki et al., 2015). In addition, it was demonstrated recently that GIRDIN acts as an effector of AMP-activated protein kinase (AMPK) under energetic stress to maintain tight junction function (Aznar et al., 2016b). Of note, these two functions are not shared by fly Girdin (Garcia-Marcos et al., 2009; Ghosh, 2017), and were thus acquired by GIRDIN during evolution to fulfill specialized functions. In contrast, our discovery of the Girdin-dependent inhibition of aPKC reveals a core mechanism contributing to epithelial cell polarization from flies to humans.

GIRDIN is considered to be an interesting target in cancer due to its role in cell motility, and high levels of GIRDIN have been reported to correlate with a poor prognosis in some human cancers [(Garcia-Marcos et al., 2015; Weng et al., 2010), and not shown]. Notwithstanding that GIRDIN may favor tumor cell migration, our study indicates that inhibition of GIRDIN function in the context of cancer would be a double-edged sword for many reasons. Indeed, we showed that knockdown of *GIRDIN* exacerbates the impact of aPKC overexpression, and leads to overgrowth and lumen filling of Caco-2 cell cysts. Of note, overexpression of aPKC can lead to cell transformation, and was associated with a poor outcome in several epithelial cancers (Archibald et al., 2015; Parker et al., 2014). Our study thus establishes that inhibiting GIRDIN in patients showing increased aPKC expression levels could worsen their prognosis. According to our data, abolishing GIRDIN function in tumor cells with decreased levels of the human Lgl protein LLGL1, as reported in many cancers (Lin et al., 2015), could also support the progression of the disease by altering the polarity phenotype. We observed cell detachment and dissemination from *GIRDIN* knockdown cysts, thus showing that GIRDIN is required for the cohesion of multicellular epithelial structures. Of note, cells, either individually or as clusters, detaching from cysts are alive and some of them remain viable. This is analogous to what was reported in *Girdin* mutant *Drosophila* embryos in which cell cysts detach from the ectoderm and survive outside of it (Houssin et al., 2015). Other phenotypes in *Girdin* mutant embryos are consistent with a role for Girdin in epithelial tissue cohesion, including rupture of the ventral midline and fragmentation of the dorsal trunk of the trachea. Mechanistically, Girdin strengthens cell–cell adhesion by promoting the association of core adherens junction components with the actin cytoskeleton (Houssin et al., 2015). A recent study established that this molecular function is evolutionarily conserved, and that GIRDIN favors the association of β-CATENIN with F-ACTIN (Wang et al., 2018). Since knockdown of *GIRDIN* results in cell dispersion from Caco-2 cell cysts, and since weakening of E-CADHERIN-mediated cell–cell adhesion contributes to cancer cell dissemination and metastasis (Jeanes et al., 2008), it is plausible that reduced GIRDIN expression contribute to the formation of secondary tumors and cancer progression. This may explain why we found that low expression levels of *GIRDIN* correlates with decreased survival in more aggressive breast cancer subtypes and lung adenocarcinoma.

In conclusion, using a sophisticated experimental scheme combining *in vivo* approaches in *D. melanogaster* with 3D culture of human cells, we defined a conserved core mechanism of epithelial cell polarity regulation. Specifically, we showed that Girdin represses the activity of aPKC to support the function of Lgl and Yrt, and ensure stability of the lateral domain. This is of broad interest in cell biology, as proper epithelial cell polarization is crucial for the morphogenesis and physiology of most organs (Tepass, 2012). In addition, the maintenance of a polarized epithelial architecture is crucial to prevent various pathological conditions such as cancer progression (Halaoui and McCaffrey, 2015). Importantly, we show that normal Girdin function putatively impairs the progression of epithelial cancers by preserving cell polarity whilst restricting cell growth and cell dissemination. Thus, our results place a caveat on the idea that GIRDIN could be an interesting target to limit cancer cell migration, and indicate that inhibition of GIRDIN in the context of cancer could be precarious. Potential drugs targeting GIRDIN would thus be usable only in the context of precision medicine where a careful analysis of aPKC, LLGL1, and E-CAD expression, as well as the polarity status of tumor cells would be analyzed prior to treatment. Inhibition of GIRDIN in patients carrying tumors with altered expression of these proteins would likely worsen the prognosis.

## Materials and methods

### *Drosophila* genetics

The following mutant alleles and transgenic lines were used in this study: *Girdin^2^* (Houssin et al., 2015), *lgl^4^* (Gateff and Schneiderman, 1974), *yrt^75^* (Laprise et al., 2006), UAS-GFP.nls (Bloomington *Drosophila* Stock Center [BDSC] Stock No. 4776), and UAS-FLAG-Girdin (Houssin et al., 2015). Germ line clone females were produced using the FLP-DFS technique (Chou and Perrimon, 1996), and were used to produce maternal and zygotic (M/Z) mutant embryos. Maternal knockdown of *aPKC* and *lgl* was achieved by crossing the driver line matαtub67;15 (obtained from D. St-Johnston, University of Cambridge, Cambridge, UK) to a line containing an inducible shRNA directed against *aPKC* (BDSC Stock No. 34332) or *lgl* (BDSC Stock No. 35773) at 25°C.

### Cell culture

Human intestinal epithelial cell line Caco-2 cells were purchased from American Type Culture Collection (ATCC). Caco-2 cells were cultured at 37°C in 5% CO_2_ in DMEM (Wisent) supplemented with 10% fetal bovine serum (Wisent), 100 U/ml penicillin (Wisent), 0.1 mg/ml streptomycin (Wisent), and 2 mM L-glutamine (Wisent). Human embryonic kidney cell line HEK293LT (ATCC) were cultured at 37°C in 5% CO_2_ in DMEM supplemented with 10% fetal bovine serum, 100 U/ml penicillin, 0.1 mg/ml streptomycin. For 3D culture, cells were seeded onto Geltrex-coated dishes at a density of 1.25 × 10^4^ cells per well and were maintained in 2% Geltrex in complete medium at 37°C under humidified atmosphere of 5% CO_2_. After 10 days in adherent culture, cells were collected for further experiments. For inhibitor studies 1.5 μM CRT-006-68-54 or water control was added to culture medium at the time of seeding cells in 3D cultures.

### Lentivirus Production and shRNA

Lentiviruses were produced by calcium phosphate transfecting HEK293LT cells in 10-cm dishes with 20 μg of lentiviral plasmid, 15 μg of packaging plasmid (psPAX2), and 6 μg of VSVG coat protein plasmid (pMD2.G). Viral supernatants were collected after 48 h, aliquoted, and frozen at –80°C. shRNAs targeting the human *GIRDIN* mRNA were obtained in pLKO.1-puro vectors from the RNAi Consortium (TRC). Caco-2 cells were infected with lentiviral supernatants and selected by the addition of 20 μg/ml puromycin. The target sequences of TRC clone number TRCN0000148551 is CCGGCTTCATTAGTTCTGCGGGAAACTCGAGTTTCCCGCAGAACTAATGAAGTTTTTTG.

### Immunofluorescence on *Drosophila* embryos

Embryos were dechorionated in 3% sodium hypochlorite for 5 min, rinsed in water, and submitted to a flash heat fixation by sequential addition of 5 ml of E-wash buffer (7% NaCl, 0.5% Triton X-100) at 80°C and 15 ml of E-wash buffer at 4°C. Embryos were then washed with PBS prior to devitellinization by strong agitation in methanol under a heptane phase (1:1), and further incubated in fresh methanol for 1 h. Saturation of non-specific binding sites was achieved by a 1-h incubation in NGT (2% normal goat serum, 0.3% Triton X-100 in PBS), which was also used to dilute primary antibodies. The following primary antibodies were used overnight at 4°C, under agitation: rat anti-Crb (Sollier et al., 2015), 1:250; mouse anti-Dlg1 [clone 4F3, Developmental Studies Hybridoma Bank (DSHB)], 1:25. Embryos were washed three times for 20 min in PBT (0.3% Triton X-100 in PBS) before and after incubation with secondary antibodies (1:400 in NGT, 1 h at room temperature), which were conjugated to Cy3 (Jackson ImmunoResearch Laboratories) or Alexa Fluor 488 (Molecular Probes). Embryos were mounted in Vectashield mounting medium (Vector Labs), and imaged with a FV1000 confocal microscope coupled to FluoView 3.0 (Olympus), using a 40× Apochromat lens with a numerical aperture of 0.90. Images were uniformly processed with Olympus FV1000 viewer (v.4.2b), ImageJ (National Institutes of Health), or Photoshop (CC 2017; Adobe).

### Cuticle preparation

Embryos were dechorionated, mounted in 100 μl of Hoyer’s mounting medium (prepared by mixing 50 ml of distilled water, 20 ml of glycerol, 30 g of gum Arabic, and 200 g of chloral hydrate)/lactic acid (1:1) and incubated overnight at 80°C. Embryos were imaged with an Eclipse 600 microscope (Nikon) through a 10× Plan Fluor objective with a numerical aperture of 0.30, and a CoolSNAP fx camera (Photometrics) coupled to MetaVue 7.77 (Molecular Devices). Images were processed with Photoshop (CC 2017; Adobe).

### Antibody production

Antibodies against the amino acids 696 to 972 of Yrt in fusion with GST were produced in rabbits (Medimabs).

### Phosphatase assays

Dechorionated embryos were homogenized in ice-cold lysis buffer (1% Triton X-100, 50 mM TRIS-HCl pH 7.5, 5% glycerol, 150 mM NaCl, 1 mM PMSF, 0.5 μg/mL aprotinin, 0.7 μg/mL pepstatin, and 0.5 μg/mL leupeptin). Lysates were cleared by centrifugation at 4°C, and 400 units of λ Phosphatase (New England Biolabs) was added to 50 μg of proteins extracted from embryos. The volume of the reaction mix was completed to 30 μl with the MetalloPhosphatase buffer (New England Biolabs) containing 1 mM of MnCl2 prior to a 30-min incubation at 30°C. The reaction was stopped by addition of Laemmli’s buffer.

### Western Blot

Dechorionated embryos were homogenized in ice-cold lysis buffer (1% Triton X-100, 50 mM TRIS HCl pH 7.5, 5% glycerol, 100 mM NaCl, 50 mM NaF, 5 mM EDTA pH 8, 40 mM β-glycerophosphate, 1 mM PMSF, 0.5 μg/mL aprotinin, 0.7 μg/mL pepstatin, 0.5 μg/mL leupeptin and 0.1 mM orthovanadate) and processed for SDS-PAGE and western blotting as previously described (Laprise et al., 2002). Primary antibodies used were: guinea pig anti-Girdin 163 (Houssin et al., 2015), 1:2,000; rabbit anti-Yrt (this study), 1:5000; mouse anti-GAPDH (Medimabs), 1:500; rabbit anti-Lgl D300 (Santa Cruz Biotechnology) 1:1,000; rabbit anti-PKCζ C20 (Santa Cruz Biotechnology), 1:2,000; mouse anti-Actin (Novus Biologicals), 1:10,00; mouse anti-Tubulin (DM1A, Sigma), 1:10,000; rabbit anti-GIRDIN (ABT80, Millipore), 1:1,000. HRP-conjugated secondary antibodies were used at a 1:2,000 to 1:10,000 dilution.

### Immunofluorescence on human 3D cysts

Three-dimensional cysts were transfected with plasmids and fixed with 2% paraformaldehyde/PBS for 10 min and permeabilized in PBS supplemented with 0.5% Triton X-100/10% goat serum for 1 h, and incubated overnight in primary antibodies. The following primary antibodies were used: mouse anti-PAR6 (1:100, Santa Cruz Biotechnology); rabbit anti-E-cadherin (1:100, Cell Signaling Technology). Cysts were washed three times for 15 min in 0.5% Triton X-100/PBS before and after incubation with secondary antibodies. Proteins were visualized with 647 Phalloidin (1:100, Invitrogen), Cy3-Donkey anti-Rabbit (1:750, Jackson IR through Cedarlane), Alexa Fluor 647-Donkey anti-Rabbit (1:200, Jackson IR through Cedarlane). DNA was detected with Hoechst dye 33258 (Sigma Aldrich). Cysts were imaged with ZEISS LSM700 confocal microscope at Plan-Apochromat 20X/0.8 M27 objective lens. Images were uniformly processed with ImageJ (National Institutes of Health).

### Cell Survival Assays

CaAM (calcein AM)/EthD-I (ethidium homodimer I) staining was performed in three-dimensional cysts with 2 mM calcein AM and 4 mM ethidium homodimer I (Live/Dead^®^ Viability/Cytotoxicity kit; Life Technologies) for 40 min at 37°C in the dark. DNA was detected with Hoechst dye 33312 (Invitrogen).

### Live imaging

Three-dimensional cysts were imaged using ZEISS LSM700 confocal microscope at 20X/0.4 Korr M27 objective lens. Cells were imaged every 20 min for 26 h in a humidified chamber with 5% CO_2_ and heated to 37°C.

### Patient cancer survival

Survival data was retrieved from the kmplot resource (kmplot.com) for breast, lung, ovarian, and gastric mRNA expression. Jetset optimal probes were selected for analysis. The breast cancer intrinsic subtypes were selected for Luminal A, Luminal B, HER2-positive, and basal cancers. Lung cancer subtypes were selected using the histology selection option for adenocarcinoma and squamous lung cancers. Survival plots with were generated using JMP14 statistical software.

### Statistical analysis

Pairwise statistics were performed using Student’s tests. Multiple comparisons were compared using ANOVA, with Tukey HSD tests. Distributions were examined using a A chi-square goodness of fit test. Differences in survival were determined using a Log-Rank test. Statistical analyses were performed with JMP14 statistical software, with α = 0.05 for all tests.

## Acknowledgments

The authors would like to acknowledge D. Bilder, U. Tepass, D. St-Johnston, the Bloomington *Drosophila* Stock Center, the *Drosophila* Genomics Resource Center, and the Developmental Studies Hybridoma Bank for reagents. Flybase was used as an important database for this work. This work was supported by operating grants from the Canadian Institute of Health Research (CIHR) to P.L. (MOP-142236) and L.M. (PJT-156271). P.L. and L.M. are Fonds de la Recherche du Quebec – Santé Research Scholars. The work was also supported by studentships from the Karassik Family Foundation (L.T-W) and Defi Canderel (R.C.).

## Author contributions

P.L., E.H., and L.M. conceived the project. C.B., L.-T.W., M. S., C.G., A.J., and R.C. performed experiments, prepared reagents, and contributed to data analysis. P.L., C.B., M.S., L.-T.W., and L. M. interpreted most results. P.L. wrote the manuscript, which was proofread by L.M. and co-authors. Finally, P.L., C.B., M.S., L.-T.W., C.G., A.J., R.C. and L. M. assembled the figures. All authors read and approved the final manuscript.

---

**Supplemental Figure 1.**
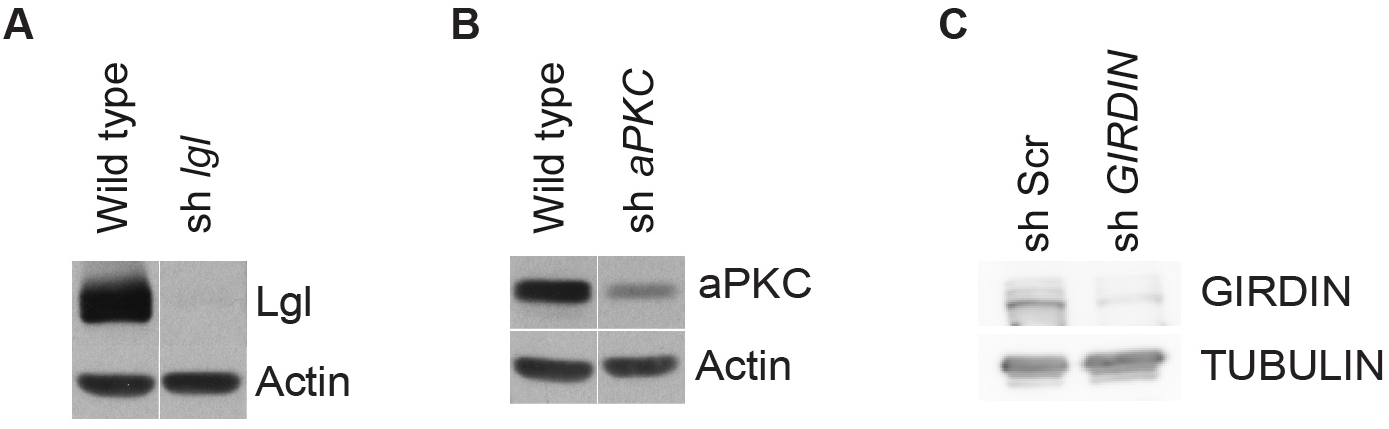

